# Transpiling reproducibility: Multi-language implementation for the St. Nicolas House Analysis algorithm using Fusion

**DOI:** 10.64898/2026.11.20.768087

**Authors:** Christopher Olvera, Alba De Prada Hernandez, Roberto Zarazua Navarro, Harshini Praveen Kumar, Piotr Fusik, Detlef Groth

## Abstract

The integration of tools written in different programming languages is common in bioinformatics pipelines. When extending these tools, developers are required to either manually port the code or use wrapper libraries, which incurs a substantial “translation tax” on developer time. Transpilers have emerged as a potential solution, enabling the generation of algorithm implementations in multiple languages from a single source code base. However, it remains essential to establish whether transpiled code preserves both functional correctness and stochastic reproducibility across language ecosystems. In this case study, we investigated whether the Fusion language and its `fut` transpiler could produce functionally identical, stochastically reproducible, and high-performing code for the St. Nicolas House Analysis (SNHA) algorithm across multiple target languages. The SNHA algorithm was implemented once in Fusion and transpiled to nine target programming languages. The generated code was evaluated against reference R, Python, and Julia implementations using the clinical `birthwt` dataset and a 26-variable synthetic benchmark dataset evaluated over 1,000 bootstrap iterations. All nine transpiled outputs produced identical network structures to the original R implementation. With the deterministic Xorshift*64 pseudo-random number generator integrated into the algorithm, all languages generated identical bootstrapped datasets and adjacency matrices. Runtimes and memory usage spanned more than two orders of magnitude: the transpiled Python implementation achieved an 8.5-fold speedup and an 8.8-fold reduction in peak memory relative to reference Python, while compiled targets (C, C++) achieved speedups exceeding 160-fold and memory reductions up to 36-fold. These findings demonstrate that modern transpilers can eliminate cross-ecosystem translation tax while guaranteeing cross-language reproducibility in computational biology.

## 1 Introduction

With the wide range of programming languages and software ecosystems available to users, developers often have considerable freedom in the selection of the language in which new tools are implemented. In contrast, this flexibility can create challenges for end-users who are trying to integrate these tools into existing bioinformatics workflows. Pipelines in bioinformatics often combine multiple tools for tasks such as genomic data processing, statistical analysis, and machine learning, where each tool may be written in a different programming language. For example, a researcher working primarily in Python may need to incorporate an algorithm from an R-based package for a particular analysis. Rather than being able to directly integrate the desired functionality, the user might need to identify an existing wrapper, manually translate the relevant source code, or implement the algorithm in Python.

For developers, the situation is equally cumbersome. Extending the reach of a biological tool by porting it to other languages (e.g., Python, R, C++) is time consuming, as each language possesses its own syntax, memory model, and optimization nuances. Furthermore, if a developer wishes to increase performance by rewriting a module in a compiled language like C, they must navigate additional complexities such as manual memory management. This “translation tax” diverts time away from other goals for both developers and users.

A modern and tempting approach to multi-language translation is the use of Large Language Models (LLMs) to translate the original implementation of the algorithm or to create equivalent code in the desired language. However, due to the probabilistic nature of these models, manual intervention is invariably required to check, debug, and reverify the output. These quality control measures incur significant costs in terms of time invested for the developer and monetary cost in computational tokens. A clear disadvantage of using these models is that the code generated with the models can violate licenses, since the code fragments they reproduce come from different parts of the internet [1].

An alternative solution lies in transpilers, motivated by the approach of automatically translating computational code between programming languages while preserving the behavior of the original implementation. A transpiler (or source-to-source compiler) transforms programs written in a source language into a target language by mapping syntax constructs to the most natural equivalent in the target environment. Unlike traditional compilers that produce machine code, a transpiler’s output must still be compiled or interpreted in the target environment. Several transpilers have been developed to facilitate inter-language interoperability. Table 1 summarizes the target languages supported by several commonly used transpilers.

**Table 1.** Transpilers, supported target languages, and latest stable releases.

| **Transpiler** | **Supported Languages** | **Release** | **Date** |
|---|---|---|---|
| Fusion (fut) [2] | C, C++, C#, D, Java, JavaScript, TypeScript, Python, Swift, OpenCL C | 3.3.8 | Sep. 21, 2026 |
| Haxe [3] | C++, C#, Java, JavaScript, JVM, Python, PHP, Lua | 4.3.7 | May 9, 2025 |
| Babel [4] | JavaScript, TypeScript | 8.0.3 | Jun. 27, 2026 |
| Temper [5] | C#, Java, JavaScript, TypeScript, Python, Lua, Rust | 0.6.0 | Dec. 30, 2025 |

Here we focus on the Fusion programming language and its transpiler `fut`, which is explicitly designed for multi-language library development from a single source code base [2]. Using Fusion, the algorithm needs to be written only once, and developers can then transpile the source code to the desired target programming language. This approach ensures that if a bug is found or a feature is extended, the correction is applied to a single source file and propagated to all targets, guaranteeing that computational logic remains consistent across the generated code.

To demonstrate the usage of this approach, we re-implemented the St. Nicolas House Analysis (SNHA) algorithm alongside its Monte Carlo graph simulation module (`mcgraph`) [6, 7]. This allowed us to evaluate whether a transpiler can achieve reproducible deterministic results across multiple languages while maintaining competitive performance. Originally, the algorithm was developed in the statistical programming language R, which limited users to exploring the tool only through use of the Comprehensive R Archive Network package [8].

To extend the reach of the algorithm to users, in the past it was ported into the programming languages Python and Julia [9–12]. Here, we see a redundant theme of reproducing the same code in different languages and more development time for each project. Through the use of the Fusion programming language transpiler, we were able to reduce some of the “translation tax” that comes with developing bioinformatics software by targeting nine of the most popular programming languages supported by the transpiler itself, and by using a wrapper to introduce more programming languages. In Fig 1, we can see an example of how an algorithm written in Fusion can be transpiled to many other languages.

**Fig 1.**
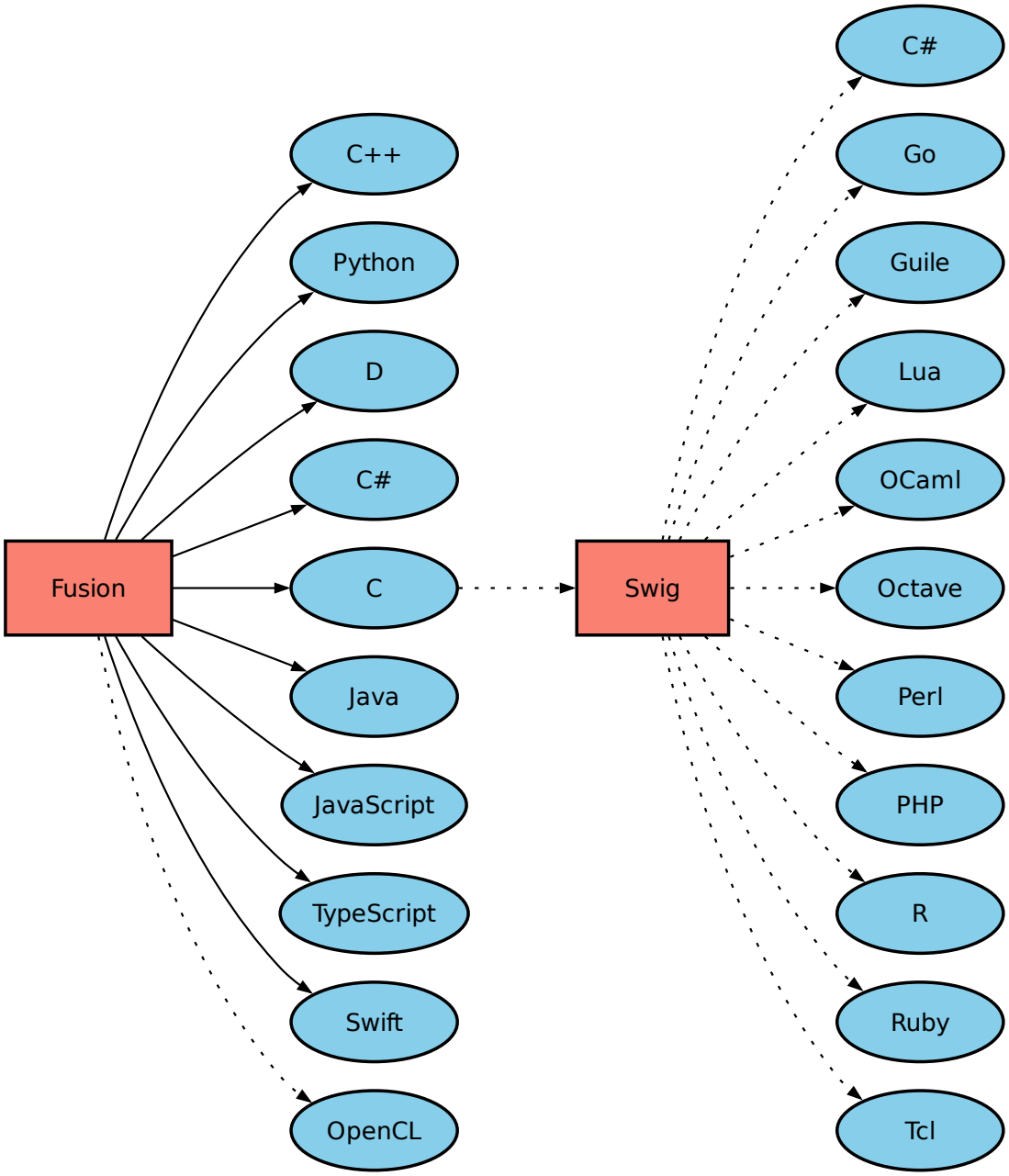
Overview of the Fusion programming language targets. The `fut` transpiler targets nine languages directly, with the option to extend coverage via SWIG wrappers around the generated C code [13]. The path from C to SWIG and from SWIG to other languages is not yet implemented, but planned for the future.

**Fig 2.**
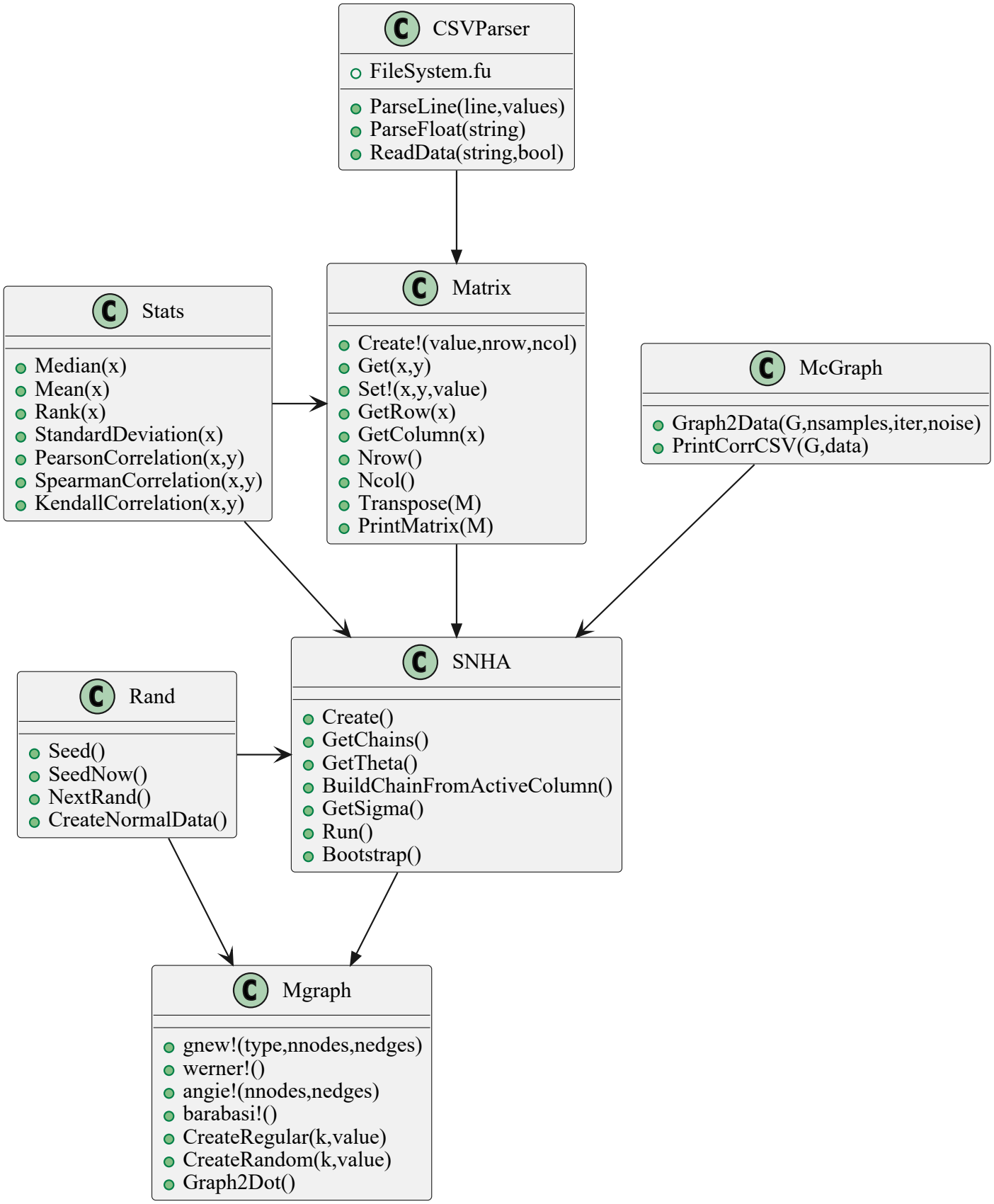
UML diagram of the Fusion classes of the project snha4any. Fig 2. UML class diagram of the snha4any **project written in Fusion**. Because Fusion transpiles a single, dependency-free codebase into multiple target languages without a dedicated runtime, the project is organized into self-contained classes to support cross-language code generation. The diagram illustrates the modular architecture and the relationships among these core classes. [19] [20]

**Fig 3.**
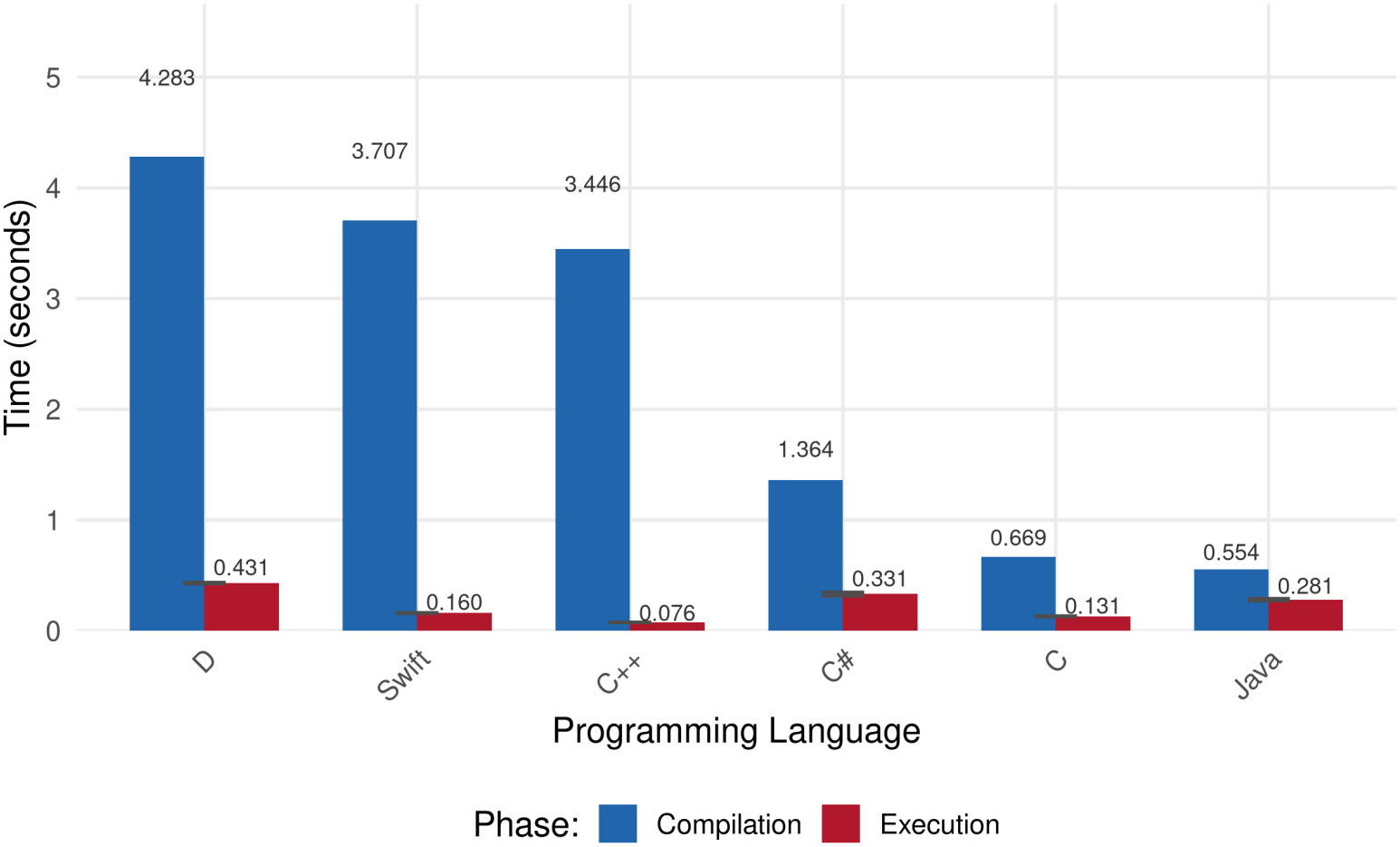
Compilation and execution times for compiled languages. For each compiled language target (C, C++, C#, D, Java, Swift), total elapsed time was separated into native compilation (mean of five runs) and execution across 1,000 bootstrap iterations (mean of ten runs). Error bars represent standard deviations.

## 2 Materials

### 2.1 The Fusion language and the fut transpiler

Fusion is an object-oriented programming language with C#-like syntax and C++-like semantics, designed for building reusable libraries that transpile into multiple target languages. It offers built-in support for mathematical functions and sophisticated data structures while remaining lightweight and self-contained. The language is self-hosted, and its `fut` transpiler currently targets C, C++, C#, D, Java, JavaScript, TypeScript, Python, Swift, and OpenCL C. Because `fut` outputs standard C, the generated code can also be bridged to additional languages,such as Tcl, R, Lua, and Octave, through interface generators such as SWIG [13]. While Fusion abstracts away much of the complexity of individual target languages, developers must still account for target-specific constraints.

### 2.2 Algorithm Overview and Experimental Design

To evaluate whether a transpiler can reliably produce reproducible, high-performance code across multiple programming languages, we re-implemented the Xorshift*64 algorithm, the St. Nicolas House Analysis (SNHA) algorithm, and its companion synthetic data generator (`mcgraph`) using the Fusion programming language. Our experimental design was structured around three core questions important to computational biologists: (1) Does the transpiled code produce functionally identical outputs across languages? (2) Does it preserve stochastic reproducibility? (3) What is the computational cost/benefit of transpilation?

### 2.3 Stochastic Reproducibility: Xorshift*64

A critical challenge in multi-language portability is ensuring that stochastic processes such as bootstrapping and Monte Carlo simulations yield identical results across different environments. Default random number generators in programming languages such as Python, R, and C use differing algorithms that result in divergent sequences even when initialized with the same seed. For example, Python’s `random` module employs the Mersenne Twister (MT19937) as its core generator, and R’s default RNG is also the Mersenne Twister [14, 15]. Despite the R version sharing the same underlying algorithm as Python, it produces different random sequences due to differing seeding mechanisms and internal state management [16]. This difference means that the same stochastic algorithm, when run across different languages or even different compilers, may produce divergent results.

To address this issue, Piotr Fusik, author of the Fusion programming language, implemented the Xorshift*64 random number generator as a module complementary to this project. Xorshift is a deterministic number generator that produces a fixed sequence of values for a given initial seed [17]. The Fusion implementation uses the same algorithm stored as a module to be utilized across all transpiled implementations. This approach provides a common set of pseudorandom numbers and removes differences arising from language-specific random number generators.

For validation, the same initial seed (123) was used in each target language for reproducible results and comparison of adjacency matrices and association chains. Bootstrapping was our main test since random number selection is used for resampling with replacement of data matrices. The resulting sequences were compared across all target languages to determine whether the generator produced identical values independently of the programming language.

### 2.4 The mcgraph Module: Random Data Generation

To test portability, we utilized the `mcgraph` module, a Monte Carlo-based synthetic data generator developed alongside SNHA in the Fusion programming language. The module takes a user-defined graph topology (e.g., band, circle, or Werner graphs) and simulates a data set via iterative propagation of node values with added Gaussian noise [7]. Because the true underlying topology is known, we can verify that the correlation matrices produced by the transpiled code faithfully reflect the input graph. This provides a quantitative benchmark for functional correctness: if the transpiled Python, C, and C++ outputs produce the same synthetic correlation matrices given the same random seed, we have demonstrated algorithmic reproducibility.

### 2.5 The St. Nicolas House Analysis Algorithm: Detecting Dependency Structures in Multivariate Data

The St. Nicolas House Analysis (SNHA) is a non-parametric, exploratory method that uncovers dependency structures among interacting variables without imposing prior assumptions about which variables are inputs or outcomes [6, 18]. A key advantage is that SNHA operates directly on association matrices, such as correlation or covariance matrices, eliminating the need for raw data. SNHA identifies dependencies by constructing association chains for each variable of interest. Beginning with a focal variable *X*_*i*_, the algorithm ranks all other variables by the magnitude of their absolute correlations with *X*_*i*_. The variable with the largest correlation becomes the first-order neighbor; continuing on, the procedure identifies the next variable with the largest correlation to that neighbor, excluding already-included variables, and continues iteratively. The chain terminates when correlation magnitude falls below a specified threshold, when correlations become insignificant, or when adding another variable would create a repetition [6]. SNHA incorporates a validation criterion to confirm that discovered chains reflect genuine dependencies rather than spurious patterns. A chain qualifies as valid only if it maintains consistent ordering when reversed. For a chain *A*-*B*-*C*-*D*, this means that starting from *A* requires |*r*_*AB*_| *>* |*r*_*AC*_| *>* |*r*_*AD*_|, while starting from *D* requires |*r*_*DC*_| *>* |*r*_*DB*_| *>* |*r*_*DA*_|. This symmetry property ensures that the dependency structure is internally coherent [6]. In addition to linear chains, SNHA identifies two other structural patterns: association bridges, in which a variable occupies a central position within a chain, and association switches, which emerge as secondary structures when end variables are removed from primary chains [18]. These elements are then synthesized into a network representation where nodes denote variables and edges represent identified dependencies. While SNHA does not inherently determine causal direction, direction can be imposed when prior knowledge of temporal or logical ordering exists, for instance when parental education precedes child developmental outcomes [18].

### 2.6 SNHA package import to Fusion Transpiler

Since the Fusion programming language was designed to be used as a library-building language, modules were built based on the fundamentals of the St. Nicolas House Analysis algorithm so that they can be imported into a programming language of choice. The following modules were created to facilitate seamless transpilation and ensure consistency across programming environments:

- **CSVParser.fu** Developed to parse comma-separated value (CSV) files, as Fusion does not provide a native library for this functionality.
- **Matrix.fu** Provides matrix data structures and numerical operations required by the statistical and network modules.
- **Rand.fu** Implements the Xorshift*64 pseudorandom number generator, which serves as the foundation for seeding and random number generation across all stochastic components of the SNHA algorithm pipeline, most importantly bootstrapping and Monte Carlo simulations.
- **Stats.fu** Provides essential statistical functions that are not native in Fusion.
- **Mgraph.fu** Handles graph generation and network visualization. Since Fusion lacks native graphics libraries, this module exports graph structures in DOT format to be rendered externally using GraphViz.
- **Mcgraph.fu** Extends graph functionality to generate synthetic datasets via Monte Carlo propagation.
- **SNHA.fu**

Contains the core implementation of the SNHA algorithm.

All generated files were fused into a single source code file and then compiled or interpreted, except for Java, where each class has to be placed in its own file.

### 2.7 Experimental Design

To evaluate the Fusion transpiler’s ability to produce correct and high-performing multi-language implementations, we designed an experimental framework that included three parts. In the first stage, we validated functional correctness using a real-world dataset. Second, we benchmarked runtime performance and memory usage using synthetic data with previously known properties. Finally, we developed a standardized benchmark protocol to ensure fair and reproducible comparisons across all target languages.

### 2.8 Functional Validation with Real-World Data

To confirm that the transpiled algorithm produces biologically meaningful and identical outputs, we analyzed the `birthwt` dataset from the `MASS` R package (Baystate Medical Center, 1986) [21]. This dataset contains risk factors associated with low infant birth weight. For each target language, we compared the SNHA association chains generated by the transpiled output against those generated by the original R implementation. Identical chain structures, in which the same ordering of variables such as (A-B-C-D) is defined, would indicate that the transpiler preserves the algorithm’s logic throughout every programming language.

### 2.9 Performance Benchmarking with Synthetic Data

To evaluate runtime performance, we generated a synthetic dataset using `mcgraph` comprising of 26 variables (nodes) and 100 samples per variable, representative of a typical moderate-scale biological dataset. The graph topology was based on a Werner graph that ensured reproducibility of the same network structure using the SNHA algorithm [7]. We initialized the SNHA algorithm across all transpiled languages using identical parameters:

- **Correlation method**: Pearson correlation coefficient
- **Chain initiation threshold**: |*r*| *>* 0.1 (minimum absolute correlation required to start an association chain)
- **Bootstrap replicates**: 1,000 (to estimate empirical edge probabilities)
- **Probability threshold**: *p >* 0.05 for edge acceptance

Bootstrapping was enabled specifically to stress-test the computational overhead of the transpiled code, as this stochastic resampling procedure is typically where performance bottlenecks emerge. The results were compared with the original Python and R implementations to quantify any performance overhead.

### 2.10 Benchmarking Protocol

To ensure fair benchmarking conditions for performance comparisons across all nine target languages, we established a unified benchmarking protocol. The following metrics were collected for each language, including the already implemented versions in R, Julia, and Python.

1. **Compilation time (compiled languages only):** The time required for the target language’s compiler to compile the Fusion-generated source code into native machine code or bytecode.
2. **Execution time**: The wall-clock runtime of the SNHA algorithm, measured from the start of the process running.
3. **Peak memory usage**: The maximum resident set size (RSS) during execution.

All benchmark metrics were automated via the Make framework to automate and guarantee identical run conditions across all language targets. The framework enforced the same synthetic input dataset (26 variables, 100 samples), identical algorithm parameters, and consistent hardware environment throughout the process. Each language implementation was executed 10 times and the median execution time was reported to mitigate variability from other processes. Unless otherwise noted, all benchmarks ran on a single CPU core; no explicit parallelization was used.

### 2.11 Timing Methodology

All timing measurements were captured using the Unix `date` command with the `+%s.%N` format, providing precise timing. Elapsed durations were computed using bc -l for high-precision arithmetic. We use this timing method as a representation of how long a user would expect to wait for the program to execute. For compiled languages (C, C++, C#, D, Java, and Swift), the native compilation step for each language was timed separately from the execution step. Each compilation was repeated five times per language to account for file system caching and potential system load variability.

Transpilation time was excluded from the timings, as the code generation step is performed once and is independent from the timings of the target language. Our benchmarks were designed to reflect only the performance of the transpiled code rather than the transpilation process itself.

For interpreted languages (Python, JavaScript, and TypeScript), no compilation step is needed; therefore, run time was measured from initiation of each program to completion. The median execution time across 10 independent runs was reported for each language to reduce the influence of sporadic system events.

### 2.12 Memory Management

Peak memory consumption was measured using the GNU `time` utility (`/usr/bin/time -v`), which reports the maximum resident set size (RSS) during the execution process. For each language, we captured the peak RSS across the 10 execution runs used for timing. The average peak memory usage, measured in megabytes (MB), was calculated to provide an estimate of expected memory consumption for each target language. All memory measurements were performed on the same hardware and under identical load conditions.

### 2.13 Hardware and Software

An overview of the compilers, runtimes, and execution environments used for performance benchmarking is given in Table 2. We tested a total of nine target languages generated from the Fusion source code: compiled (C, C++, D, Swift), just-in-time compiled (C#, Java, JavaScript,TypeScript), and purely interpreted (Python). Although Fusion supports one extra target OpenCL C, we did not benchmark this GPU-oriented language in this study.

**Table 2.** Compilers, runtimes, and optimization configurations used for performance benchmarking.

| **Language** | **Compiler / Runtime** | **Optimization / Build Configuration** |
|---|---|---|
| Fusion (fut) | fut 3.3.6 | N/A |
| C | GCC 14.2.0 | -O3; -lm; ‘pkg-config --libs glib-2.0‘ |
| C++ | GCC 14.2.0 | -O3 -std=c++20 |
| C# | .NET SDK 10.0.300 | dotnet build -c Release |
| D | LDC 1.40.0 | -O |
| Java | OpenJDK 21.0.11 | N/A |
| JavaScript (Node.js) | Node.js 24.18.0 | N/A |
| JavaScript (Bun) | Bun 1.4.0 | N/A |
| TypeScript | Bun 1.4.0 | N/A |
| Python (fut) | CPython 3.13.5 | N/A |
| Swift | SwiftC 6.3.3 | swift build -c release |
| R (Groth) | Rscript 4.5.0 | N/A |
| Python (Hake) | CPython 3.13.5 | N/A |
| Julia (Zarazua) | Julia 1.13.1 | Default JIT execution (julia SNHA_Test.jl) |

All performance tests were conducted on a single machine to eliminate hardware variability. The system was an x86 64 architecture with an 11th Gen Intel Core i7-11800H processor (16 logical cores, 2.30 GHz base frequency) and 16 GB of RAM, running Debian GNU/Linux 13 (trixie) with kernel 6.12.96. The CPU frequency governor was set to `performance` during benchmarking.

For compiled languages, we used aggressive optimization flags where available: GCC with -O3 for C and C++, LDC with -O for D, and release builds for C# and Swift. For JavaScript and TypeScript, we used both Node.js (v24.18.0) and Bun (v1.4.0) runtimes to evaluate performance differences across JavaScript and TypeScript engines. The Fusion transpiler `fut` version used for code generation was 3.3.6. All benchmarking was automated via the project Makefile to ensure reproducible execution conditions, including consistent environment variables, file paths, and output redirection.

Compilation and transpilation times were excluded from runtime measurements and only the execution time of the pre-built binaries or scripts was recorded.

## 3 Results

### 3.1 Functional Equivalence Across Target Languages

To validate whether the Fusion transpiler implementations preserve algorithmic logic, we compared the SNHA association chains generated by each transpiled language against the original R implementation using the clinical `birthwt` dataset from the `MASS` package [21].

The reference R implementation identified six association chains linking clinical risk factors for low birth weight. All nine transpiled implementations (Python, C, C++, C#, D, Java, JavaScript, TypeScript, and Swift) recovered the exact same six chains. Pearson correlation coefficients computed by each implementation agreed with the R reference within floating-point precision (*<* 10^*−*6^). Table 3 presents a comparison of the association chains generated by the transpiled languages and the R reference.

**Table 3.**
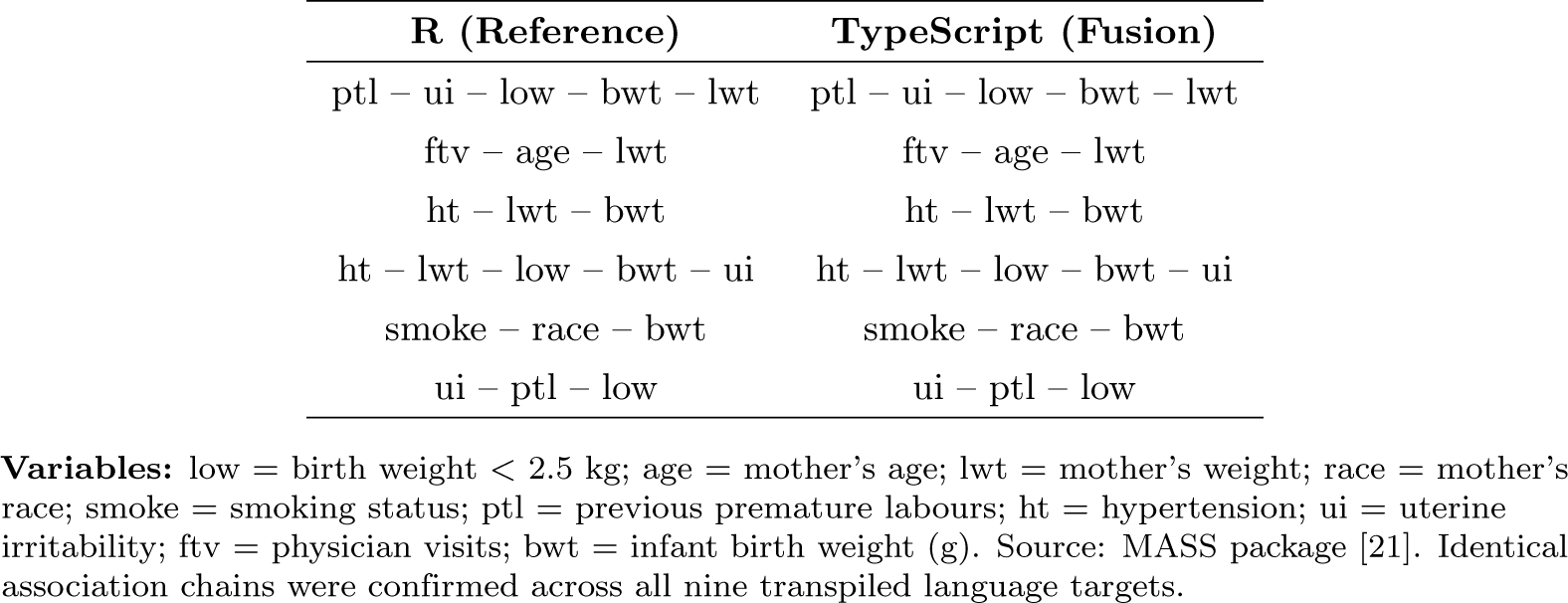
Comparison of SNHA association chains between reference R and transpiled implementations.

| **R (Reference)** | **TypeScript (Fusion)** |
|---|---|
| ptl – ui – low – bwt – lwt | ptl – ui – low – bwt – lwt |
| ftv – age – lwt | ftv – age – lwt |
| ht – lwt – bwt | ht – lwt – bwt |
| ht – lwt – low – bwt – ui | ht – lwt – low – bwt – ui |
| smoke – race – bwt | smoke – race – bwt |
| ui – ptl – low | ui – ptl – low |

**Variables:** low = birth weight < 2.5 kg; age = mother’s age; lwt = mother’s weight; race = mother’s race; smoke = smoking status; ptl = previous premature labours; ht = hypertension; ui = uterine irritability; ftv = physician visits; bwt = infant birth weight (g). Source: MASS package [21]. Identical association chains were confirmed across all nine transpiled language targets.

**Table 4.** Comprehensive benchmark summary across evaluated language targets (1,**000 bootstrap iterations, Werner synthetic dataset)**.

| **Implementation** | **Compile Time (s)** | **Run Time (s)** | **Peak RSS (MB)** |
|---|---|---|---|
| C++ (Fusion) | 0.58 ± 0.02 | 0.076 ± 0.001 | 3.81 |
| C (Fusion) | 0.42 ± 0.01 | 0.131 ± 0.003 | 3.19 |
| TypeScript (Bun) | N/A | 0.157 ± 0.003 | 49.76 |
| JavaScript (Node.js) | N/A | 0.160 ± 0.004 | 68.13 |
| Swift (Fusion) | 2.15 ± 0.05 | 0.160 ± 0.003 | 18.92 |
| Java (Fusion) | N/A | 0.281 ± 0.010 | 162.64 |
| C# (Fusion) | 1.84 ± 0.04 | 0.331 ± 0.016 | 48.36 |
| D (Fusion) | 1.12 ± 0.03 | 0.431 ± 0.005 | 104.84 |
| Python (Fusion) | N/A | 2.560 ± 0.091 | 13.18 |
| *Reference Implementations* |  |  |  |
| R (snha) | N/A | 4.272 ± 0.101 | 66.23 |
| Julia (snha4jl) | N/A | 20.477 ± 2.592 | 700.00 |
| Python (snha4py) | N/A | 21.757 ± 0.612 | 115.45 |

### 3.2 Stochastic Reproducibility: The Xorshift*64 Guarantee

To validate the stochastic reproducibility of the Xorshift*64 random number generator across all transpiled languages, we tested its behavior within the SNHA bootstrapping procedure. Bootstrapping involves resampling rows from the original dataset with replacement, where each row selection is determined by a pseudorandom number. Given the same initial seed (123) across all implementations, two outcomes were verified: the generated bootstrapped datasets were bitwise identical prior to chain construction, and the resulting adjacency matrices were identical.

We performed five independent bootstrapping runs for each transpiled language using seed 123. In every instance, the bootstrapped matrices and resulting adjacency graphs were identical across all target environments. This confirmed that the in-tree Xorshift*64 implementation produces uniform random sequences regardless of host architecture, compiler, or language runtime.

### 3.3 Runtime Performance Across Target Languages

We benchmarked execution times across all nine transpiled implementations as well as the reference baseline implementations in R [8], Python [10], and Julia [12]. Performance was evaluated over 1,000 bootstrap resampling iterations on the 26-variable synthetic Werner dataset (*N* = 100) [7]. Reported metrics represent the mean and standard deviation across ten independent runs per implementation.

Mean execution time spanned more than two orders of magnitude, ranging from 0.076 s (compiled C++) to 21.757 s (reference Python). Compiled C++ (0.076 s, SD 0.001 s) and C (0.131 s, SD 0.003 s) achieved the shortest execution runtimes, delivering 286.3-fold and 166.1-fold speedups over reference Python, and 56.2-fold and 32.6-fold speedups over reference R (4.272 s, SD 0.101 s), respectively. TypeScript on Bun (0.157 s, SD 0.003 s), JavaScript on Node.js (0.160 s, SD 0.004 s), and compiled Swift (0.160 s, SD 0.003 s) executed within a similar range to native C. Intermediate runtimes were observed for managed environments: Java (0.281 s, SD 0.010 s), C# (0.331 s, SD 0.016 s), and D (0.431 s, SD 0.005 s).

Among dynamic and interpreted environments (Fig 4), transpiled Python achieved an average runtime of 2.560 s (SD 0.091 s), representing an 8.5-fold speedup over reference Python (21.757 s, SD 0.612 s) and a 1.7-fold speedup over reference R. The reference Julia implementation recorded a mean execution time of 20.477 s (SD 2.592 s; range: 17.894–25.620 s), performing on par with reference Python.

**Fig 4.**
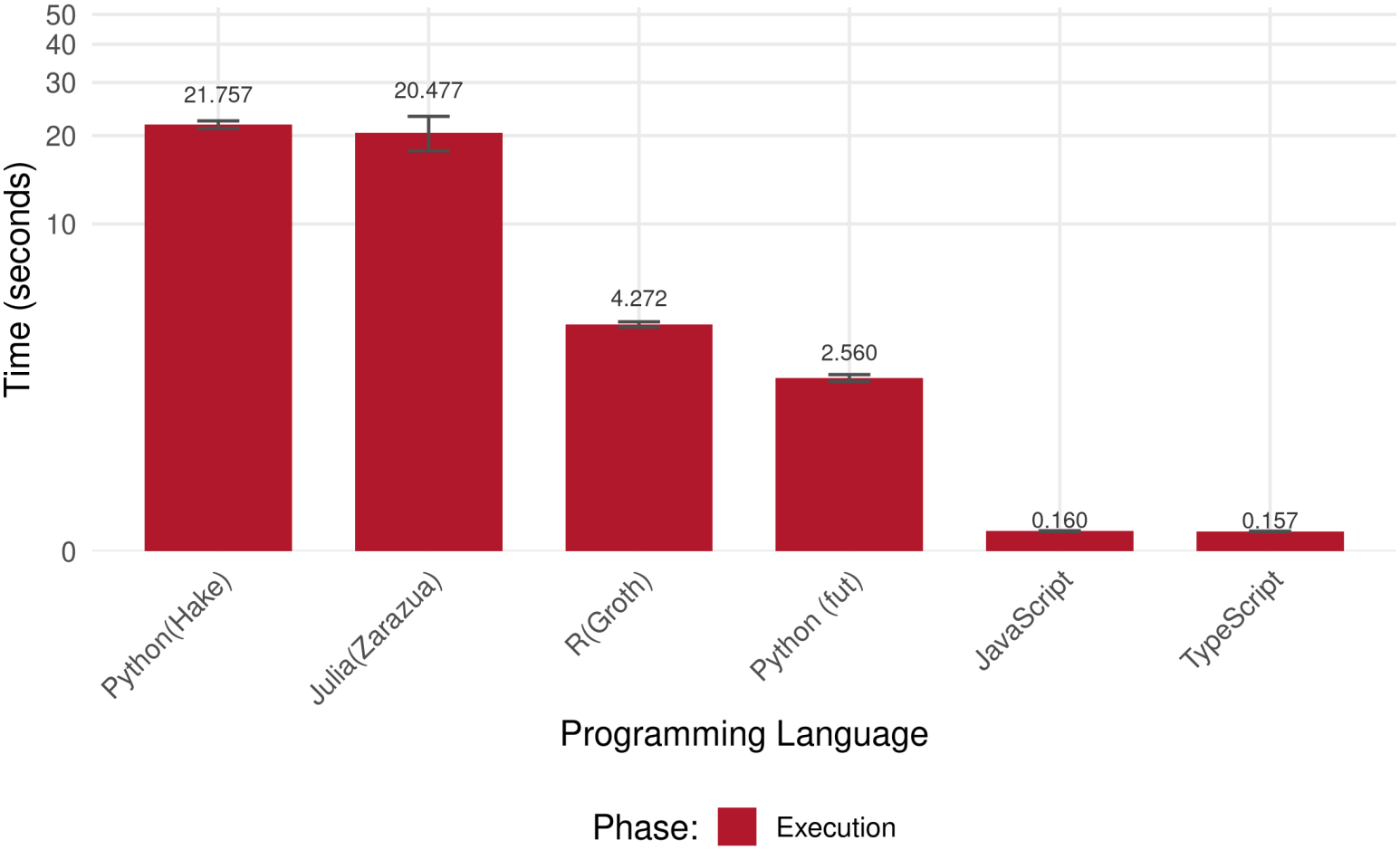
Execution times for interpreted and dynamically compiled environments. Runtimes for interpreted and JIT-compiled languages are shown on a logarithmic scale across 1,000 bootstrap iterations on the synthetic Werner dataset.

### 3.4 Memory Consumption

Peak memory consumption, measured as maximum Resident Set Size (RSS), varied by more than two orders of magnitude across environments (Fig 5). Across all evaluated implementations, peak RSS ranged from 3.19 MB (compiled C) to 700.00 MB (reference Julia).

**Fig 5.**
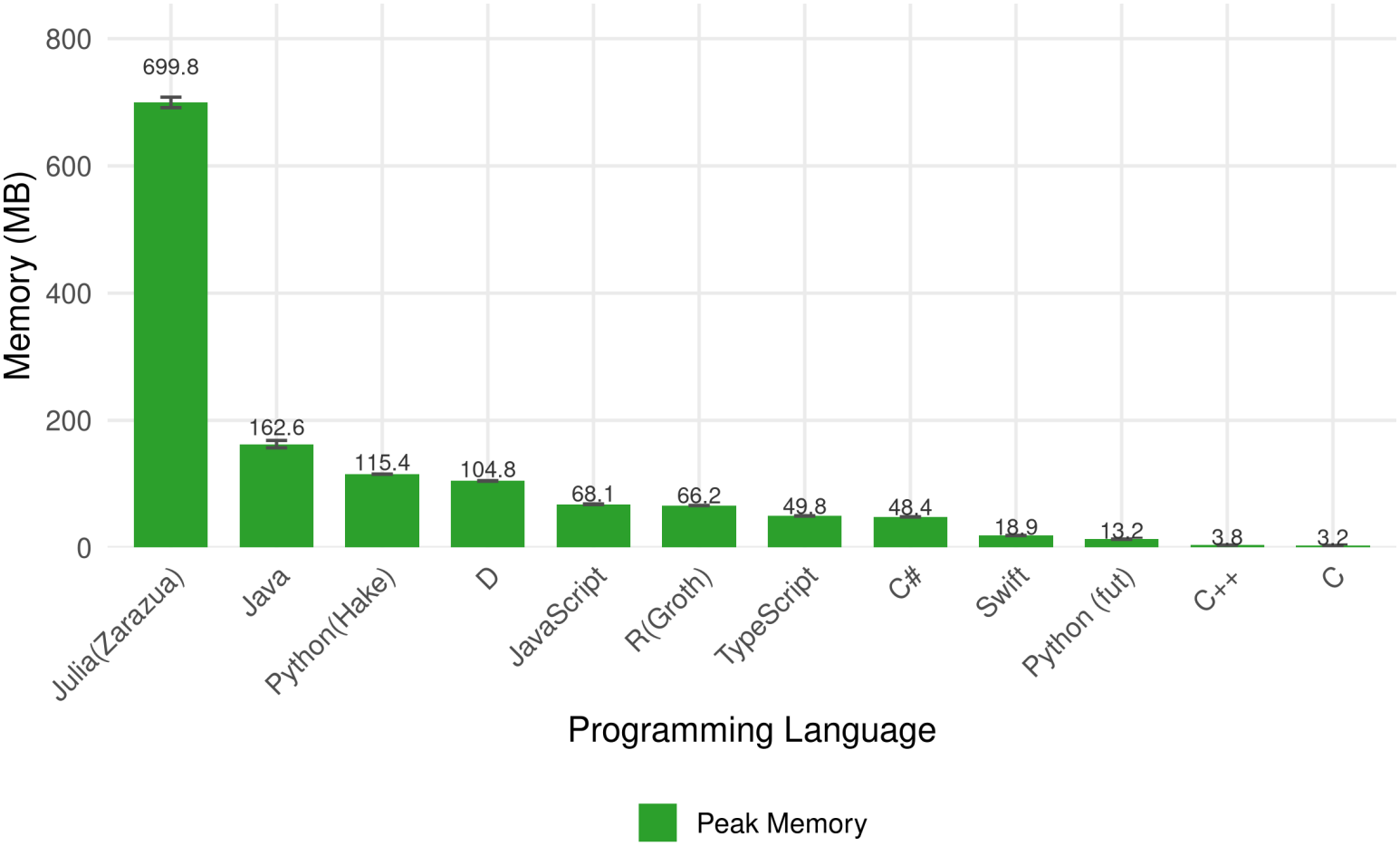
Peak memory consumption across all language implementations. Bars depict the average maximum Resident Set Size (RSS in MB) across ten independent execution runs performing 1,000 bootstrap iterations on the synthetic Werner dataset. Fusion-generated targets are compared against standalone baseline implementations in R (`snha` by Groth), Python (`snha4py` by Hake), and Julia (`snha4jl` by Zarazua Navarro). Error bars denote standard deviations.

Compiled implementations recorded the lowest memory footprints: native C required 3.19 MB, C++ consumed 3.81 MB, and Swift required 18.92 MB. Transpiled Python exhibited a peak memory usage of 13.18 MB, achieving an 8.8-fold reduction relative to reference Python (115.45 MB) and a 5.0-fold reduction relative to reference R (66.23 MB).

Managed and JIT runtime environments displayed intermediate memory footprints: C# required 48.36 MB, TypeScript on Bun consumed 49.76 MB, JavaScript on Node.js consumed 68.13 MB, D consumed 104.84 MB, and Java required 162.64 MB. Among the reference baselines, reference Julia recorded the highest peak memory consumption at 700.00 MB (SD 8.25 MB).

Average runtimes represent ten independent runs. Standalone implementations in R (`snha` by Groth), Python (`snha4py` by Hake), and Julia (`snha4jl` by Zarazua Navarro) provide baseline comparison against Fusion-generated targets (TypeScript, JavaScript, and Python).

## 4 Discussion

This work addressed a common problem in computational biology: the translation tax that comes with every new implementation of the same algorithm and the reproducibility of the same algorithm across many programming languages. With the wide range of programming languages available to users, transpilers solve this crisis of rewriting the same algorithm redundantly to reach a greater audience. This allows developers to reduce the amount of code they need to write and maintain, effectively saving time as the source code needs to be updated once and output to as many programming languages as the transpiler supports.

### 4.1 Functional and Stochastic Equivalence Across Languages

The identical recovery of all six association chains in all nine programming languages that were found in the reference R version matters biologically. Each chain, for example, ptl–ui–low–bwt–lwt, represents a hypothesized structure among clinical risk factors for low birth weight. If even one edge were to be reordered or dropped, the biological interpretation of the network would shift, regardless of whether the code otherwise ran correctly. Exact structural agreement is therefore essential across all nine languages.

The bootstrapping and final adjacency matrix check of the Xorshift*64 generator strengthens this point for the stochastic parts of the algorithm. If random number generation diverged by language, it would silently produce different bootstrapped datasets and, in effect, different adjacency matrices. This is a meaningful reproducibility guarantee for collaborative research. For example, if a collaborator were to re-run a bootstrapped SNHA analysis in another programming language using the same seed, they should get the same edge probability estimates and chains as the original author, not just a qualitatively similar network.

### 4.2 Strategic Advantages of Fusion over Alternative Transpilers

To contextualize the trade-offs of this approach, Fusion can be compared with existing source-to-source compilers such as Haxe [3] and Temper [5]. While these tools also address multi-language portability, Fusion offers several practical advantages for scientific computing and reproducible research.

First, unlike Haxe (which emits C++ coupled to the `hxcpp` runtime) or Temper (which primarily targets Rust and C#), Fusion can transpile directly to clean, portable C [2, 3, 5]. In computational biology, C remains the universal common denominator for interoperability. Standalone C source files compile natively across platforms without requiring dedicated runtime engines. Furthermore, emitting standard C simplifies integration with interface generators like SWIG [13]. Through SWIG, a single generated C codebase can be wrapped into native extensions for environments such as R, Python, Tcl, Lua, and GNU Octave, extending ecosystem reach beyond Fusion’s built-in language backends (Fig 1).

Second, Fusion resolves the problem of cross-language stochastic reproducibility. Most general-purpose transpilers rely on the host platform’s standard library for random number generation, which causes identical seeds to produce divergent random streams in Python, JavaScript, and R. By embedding the deterministic Xorshift*64 algorithm [17] directly into the Fusion source code (`Rand.fu`), the generated code produces bitwise-identical pseudorandom sequences across every target runtime. Finally, the transpiled files are self-contained and free from heavy framework dependencies, allowing compiled C and transpiled Python implementations to run within minimal memory footprints.

### 4.3 Performance and Memory Trade-Offs

The benchmark results also highlight notable efficiency differences between transpiled code and standard high-level libraries. The transpiled Python implementation ran 8.5 times faster than reference Python (`snha4py`) [10] (2.560 s vs. 21.757 s) and used nearly nine times less memory (13.18 MB vs. 115.45 MB).

The benchmark of the Julia implementation (`snha4jl`) [12] illustrates the practical trade-offs of standalone command-line scripts in dynamic environments. While Julia achieves near-C execution speeds during interactive sessions once functions are compiled, standalone command-line execution incurs compilation and package-loading latency on every invocation. In our single-run batch test, Julia clocked in at 20.477 s with a peak memory footprint of 700.00 MB, performing on par with interpreted Python. For persistent interactive sessions, this overhead is negligible; however, in high-throughput workflows (such as Nextflow or Slurm pipelines) where thousands of tasks are dispatched as independent batch processes, the startup costs compound quickly. Under those constraints, a compiled Fusion binary finishes in a fraction of a second with almost no memory overhead.

A factor to consider is that our benchmark dataset is relatively small compared with high-throughput applications. Genome-wide studies and large-scale genomic analyses can involve hundreds or thousands of variables and observations. The relative rankings of correlation coefficients determined in the SNHA algorithm were for a dataset consisting of twenty-six variables. The performance and memory gaps observed between languages might shift at larger scales, where memory bandwidth and cache locality become dominant. Nonetheless, current results show that transpiled code is highly competitive and that the Werner graph benchmark provides a rigorous basis for controlled comparison. Future work should evaluate additional network topologies to explore scaling behavior.

### 4.4 Disadvantages of Transpilers

At the time of writing, the concept of using transpilers is not new. However, adopting a relatively new programming language, such as Fusion, presents unique challenges associated with the infancy stage of any language. When this project began, Fusion lacked the extensive libraries that developers in more established languages take for granted. We found ourselves building fundamental components from scratch, components that bioinformaticians typically use on a day-to-day basis, such as CSV parsers, data structures, and statistical/mathematical libraries.

Due to these limitations at the time, we structured the project around these obstacles and developed the modules needed to make the algorithm work. We developed our matrix module to handle numerical operations, a dedicated parser for comma-separated value files, and a workaround for visualizing networks without relying on the native graphics libraries of each target programming language. Fusion does provide terminal input/output handling, but it does not provide native libraries for all domain-specific tasks; therefore, modules such as the CSV parser had to exist so that portability was maximized to support the SNHA algorithm in as many languages as possible.

The process was instructive but also time consuming. Writing code that functions and performs efficiently across multiple programming environments required careful attention to the distinct rules and constraints of each programming language. For example, C demanded explicit memory management, whereas Python handled garbage collection automatically. However, one of the advantages of Fusion is that it hides much of the complexity of the target languages: writing C-compatible code in Fusion is generally easier than writing the equivalent C code directly, and developers do not need to be experts in Swift or Java to generate working code for those targets. At the same time, some awareness of target-language constraints remains necessary to avoid transpilation or compilation problems.

Despite these obstacles, our approach and trials in this new environment led to a discovery and achievement: reproducible stochastic results. By implementing the random number generator Xorshift*64 in the transpiler source code, we ensured that the generator operated identically across all transpiled languages and produced the same results. It ensured that when running the same seed, it produced the same sequence of pseudorandom numbers in every target environment. This was essential for stochastic variability in bootstrapping and Monte Carlo simulations, which is of great importance to scientific reproducibility.

## 5 Conclusion

Our work addresses a persistent obstacle in computational biology: methods developed by bioinformaticians are frequently locked into a single language ecosystem, constraining their reach and complicating reproducibility across pipelines used in modern genomics, machine learning, transcriptomics, and network biology. Tools like Fusion allow developers to focus on the biological and statistical correctness of their algorithm rather than maintaining multiple implementations for every downstream user.

## 6 Future Work

Further exploration can be done to determine whether programming with transpilers is more efficient and time saving. First, direct benchmarking against other transpilers (Haxe, Temper) would provide a more comprehensive view of how transpilers perform and their usability. Haxe, for example, may offer a larger library infrastructure, but it does not provide C output in the same way; a balanced comparison should therefore focus on target coverage, library needs, and runtime performance rather than library count alone. Second, extending the Fusion integration of the St. Nicolas House Algorithm to its full capacity with features that are not yet present in the R and Python versions, such as additional statistical methods, would be valuable. In our tests, we benchmarked the core implementation of the algorithm with bootstrapping, but many additional functionalities are still yet to be featured, such as confusion matrices. Third, the Fusion–C–SWIG path could be used to extend the library to other languages such as Tcl, R, Python, Lua, and Octave. Finally, exploring the scalability of the St. Nicolas House Algorithm to high-dimensional datasets containing hundreds or thousands of variables, typical of modern transcriptomics, genome-wide association studies, and single-cell sequencing, remains an essential direction for future optimization.

## Supporting information

**S1 File. Source code, benchmark harnesses, and reproduction datasets**. Complete source code for the Fusion implementation `(SNHA.fu, Mcgraph.fu, Matrix.fu, CSVParser.fu, Rand.fu, Stats.fu, Mgraph.fu, Timeit.fu, FileSystem.fu, ListUtils.fu, FloatVector.fu),` target-language transpiled sources

`(cm/, cppm/, dm/, javam/, jsm/, pym/, swiftm/, tsm/),` automated benchmark execution scripts `(Makefile, SNHA Test.R, SNHA Test.py, SNHA Test.jl),` raw benchmark timing and memory measurements (bench results.csv), and reproduction datasets `(werner-synthetic.csv, birthwt.csv)` are available in the public GitHub repository: https://github.com/mittelmark/snha4any [23].

**S1 Table. Full benchmark timings, compilation times, and peak resident set size measurements**. Comprehensive summary of compilation times, execution durations, standard deviations, and peak memory consumption across all twelve evaluated language implementations over 1,000 bootstrap resampling iterations on the 26-variable synthetic Werner dataset.

## Acknowledgments

The authors thank Piotr Fusik for creating the Fusion programming language and transpiler (fut) and for implementing the deterministic Xorshift*64 pseudorandom number generator module utilized in this work.

